# Sterol-response pathways mediate alkaline survival in diverse fungi

**DOI:** 10.1101/2020.03.26.010983

**Authors:** Hannah E. Brown, Calla L. Telzrow, Joseph W. Saelens, Larissa Fernandes, J. Andrew Alspaugh

## Abstract

The ability for cells to maintain homeostasis in the presence of extracellular stress is essential for their survival. Stress adaptations are especially important for microbial pathogens to respond to rapidly changing conditions, such as those encountered during the transition from the environment to the infected host. Many fungal pathogens have acquired the ability to quickly adapt to changes in extracellular pH to promote their survival in the various micro-environments encountered during a host infection. For example, the fungal-specific Rim/Pal alkaline response pathway has been well characterized in many fungal pathogens, including *Cryptococcus neoformans*. However, alternative mechanisms for sensing and responding to host pH have yet to be extensively studied. Recent observations from a genetic screen suggest that the *C. neoformans* sterol homeostasis pathway is required for growth at elevated pH. This work explores interactions among mechanisms of membrane homeostasis, alkaline pH tolerance, and Rim pathway activation. We find that the sterol homeostasis pathway is necessary for growth in an alkaline environment, and that an elevated pH is sufficient to induce Sre1 activation. This pH-mediated activation of the Sre1 transcription factor is linked to the biosynthesis of ergosterol, but is not dependent on Rim pathway signaling, suggesting that these two pathways are responding to alkaline pH independently. Furthermore, we discover that *C. neoformans* is more susceptible to membrane-targeting antifungals in alkaline conditions highlighting the impact of micro-environmental pH on the treatment of invasive fungal infections. Together, these findings further connect membrane integrity and composition with the fungal pH response and pathogenesis.

## Introduction

Diverse cell types, from simple unicellular microorganisms to complex multicellular eukaryotes, interpret alterations in extracellular pH as a common signal for changes in the external environment. Pathogenic microorganisms are often uniquely exposed to wide fluctuations in pH as they move between various micro-environments in the human host. Among these, fungi that cause invasive fungal infections (IFIs) have acquired the ability to rapidly adapt to changes in extracellular pH to promote their survival during an infection. The shift of a fungal pathogen from an acidic external environment to the neutral/alkaline pH of the mammalian host is associated with the activation of the fungal-specific Rim/Pal signaling pathway, triggering cellular changes important for survival in these new conditions. These changes include alterations in the cell wall, often accompanied by larger morphological transitions that promote host colonization. In the common fungal pathogen *Candida albicans*, pH-directed cellular responses includes the ability to transition between yeast-like growth and invasive hyphal forms (1–3). The opportunistic human fungal pathogen and basidiomycete yeast *Cryptococcus neoformans* similarly activates Rim signaling to respond to changes in pH. Because *C. neoformans* initially colonizes the human lung, which is often relatively more alkaline than its natural environmental reservoirs, this signaling pathway is activated in the setting of infection. In fact, the *C. neoformans* Rim101 transcription factor, the terminal component of the Rim pathway, is among the most highly induced proteins *in vivo* (4).

Given its pH-dependent activation, as well as its important role in the adaptation of fungal cells to elevated pH, the Rim signaling cascade is often considered to be the major alkaline pH response pathway in fungi. However, other cellular processes and pathways are required for fungal growth under conditions of extreme pH (both acidic and alkaline). These processes include the production of glycosphingolipids (GSLs) that associate with proteins in the outer leaflet of fungal plasma membranes to form lipid rafts and maintain membrane fluidity and organization (5–7). Recent studies have demonstrated that mutations resulting in reduced or absent GSLs render yeasts such as *Kluyveromyces lactis*, *Neurospora crassa,* and *C. neoformans* unable to grow in alkaline environments (8–11). The connection between membrane composition and the ability for fungal cells to grow in alkaline environments has been associated with defects in cytokinesis, altered activity of plasma membrane proton pumps, as well as an altered lipid profile (10). Furthermore, reduced ergosterol content in membranes has been linked to salt stress sensitivity in *Saccharomyces cerevisiae* (12, 13) and to aberrant V-ATPase regulation of pH gradients in *Candida albicans* (13, 14).

Recent observations from our genetic screen suggest that *C. neoformans* sterol homeostasis might also be required for growth at elevated pH (15). The sterol homeostasis pathway (SREBP pathway) has been extensively studied in both mammalian and fungal cells (16–20). Proteins in this pathway regulate the production and delivery of sterols to the plasma membrane to maintain appropriate cell homeostasis (17, 21, 22). In several fungal species, including *C. neoformans,* the Sre1 transcription factor (the terminal transcription factor in this sterol homeostasis pathway) is activated in response to low oxygen conditions (21, 23–26). Upon activation of the *C. neoformans* sterol homeostasis pathway, the basidiomycete-specific Stp1 protease cleaves Sre1, freeing its N-terminus to release from the membrane of the endoplasmic reticulum and translocate to the nucleus (22). This cleavage is induced in an O_2_-dependent manner and is important for the transcription of many ergosterol biosynthesis genes (23, 25). However, the association between the sterol homeostasis pathway and pH adaptation has not yet been explored.

Here we define potential interactions among fungal sterol homeostasis, alkaline pH tolerance, and Rim pathway activation. We find that the sterol homeostasis pathway is indeed necessary for growth in an alkaline environment, and that an elevated pH is sufficient to induce Sre1 cleavage and activation. This pH-mediated activation of the Sre1 transcription factor is not dependent on Rim pathway signaling, suggesting that these two pathways are responding to alkaline pH independently. Furthermore, we demonstrate that Sre1-mediated ergosterol biosynthesis is linked to the response to alkaline pH and relevant in biologically diverse fungi. Finally, we discover that *C. neoformans* is more susceptible to membrane-targeting antifungals in alkaline conditions, highlighting the impact of micro-environmental pH on the treatment of this invasive fungal infection. Together, these findings connect a highly conserved pathway involved in membrane homeostasis and sterol maintenance to the adaptive response to changes in extracellular pH.

## Results

### Convergent and divergent phenotypes of the *sre1*Δ and *rim101*Δ mutants

A recent forward genetic screen identified two elements of the *C. neoformans* sterol homeostasis pathway, the Sre1 transcription factor and its activating protease Stp1, as proteins required for growth of this pathogenic fungus at an alkaline pH (15). To confirm this observation, we generated and acquired multiple, independent *C. neoformans sre1*Δ mutants, and verified that all demonstrated a severe growth defect at high pH (Fig 1A). We performed detailed phenotypic comparisons between mutants in the alkaline-responsive Rim pathway and the sterol homeostasis pathway, as exemplified by the *rim101*Δ and *sre1*Δ transcription factor mutant strains, respectively. Both mutant strains grew similarly to wildtype on a rich growth medium at pH 5.5 (YPD medium). These two mutants also displayed similar growth defects compared to wildtype on growth medium buffered to a pH greater than 7 (Fig 1A). Importantly, the *sre1*Δ alkaline pH-sensitive mutant phenotype was rescued by the reintroduction of the wildtype *SRE1* allele (Fig S1).

**Figure 1.**
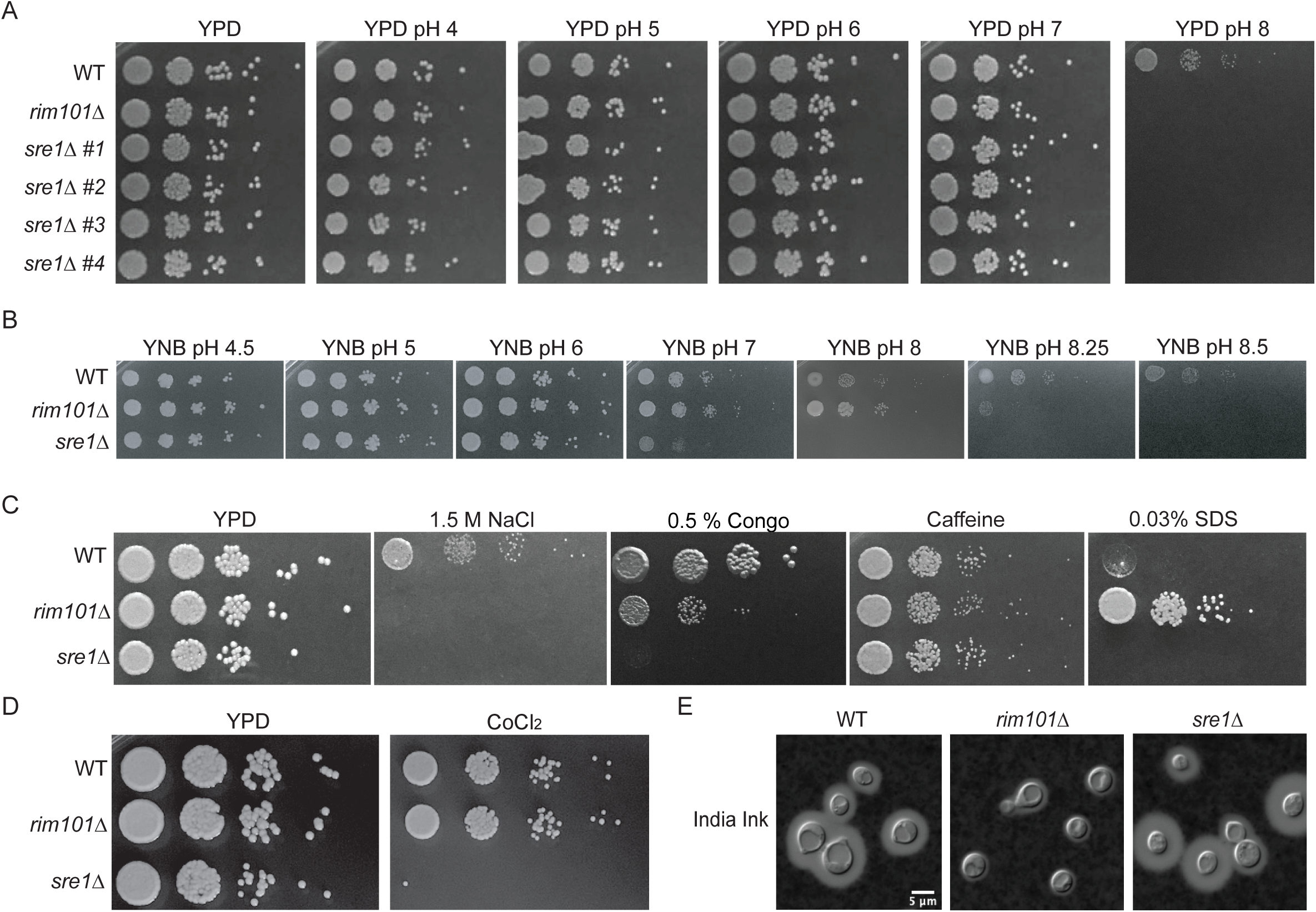
Stress response phenotypes of the *sre1*Δ and *rim101*Δ mutant strains. **A.** Four independent *sre1*Δ mutant strains were serially diluted onto YPD medium and YPD pH 4-8. Growth was compared to wildtype (WT) and a *rim101*Δ mutant known to have alkaline pH sensitivity. Growth was assessed after 3 days. *sre1*Δ #1 (HEB5) will be shown for all subsequent phenotyping and analysis. B. The *sre1*Δ and *rim101*Δ mutant strains are unable to grow at increasing pH levels on minimal media (YNB). Strains were spotted in serial dilutions onto YNB medium buffered to pH 4.5-8.5 and growth was compared to WT after 3 days. C. The *sre1*Δ and *rim101*Δ mutant strains display distinct and overlapping phenotypes to cell stressors. Strains were serially diluted and spotted to either YPD, YPD + 1.5 M NaCl, YPD + 0.5% Congo Red, YPD + 1 mg/ml caffeine, or YPD + 0.03% SDS. Growth was compared to WT and assessed after 3 days. D. The *sre1*Δ mutant strain displays a growth defect in response to hypoxia-mimicking growth conditions (7mM CoCl_2_). Strains were spotted in serial dilutions onto YPD at 30°C and YES + 7 mM CoCl_2_ at 30° C. Growth was assessed after 3 days and compared to WT and the *rim101*Δ mutant. E. The *sre1*Δ mutant strain does not have the same capsule deficiency as the *rim101*Δ mutant strain. Strains were incubated in CO_2_-independent medium for 3 days before imaging using India ink exclusion counterstaining. Capsule is noted as a halo of clearing around the yeast cells.

To account for a potential confounding effect on growth by exogenous lipids in the yeast extract-rich medium, we also assessed the ability for these mutant strains to grow on a minimal medium (YNB) buffered to a range of pH values. The *sre1*Δ and *rim101*Δ mutants were able to grow on YNB medium buffered to pH 4 through pH 7. At more alkaline pH, the growth defect of the *sre1Δ* mutant strain was more severe than that of the *rim101*Δ mutant, with the *sre1*Δ mutant unable to grow at pH 8, and the *rim101*Δ strain only displaying complete growth inhibition at pH > 8 (Fig 1B).

Given the established role of Sre1 in mediating growth in hypoxia, we compared growth rates of these mutant strains in the presence of cobalt chloride (CoCl_2_), a hypoxia-mimicking agent that disrupts many steps in the ergosterol biosynthesis pathway (27). Consistent with previous reports, (21, 24, 25, 28), the *sre1*Δ mutant is unable to grow under conditions of reduced oxygen concentration (Fig 1D and Fig S1). The *rim101*Δ mutant did not share this hypoxia-related growth defect (Fig 1D). Therefore, although sharing a similar alkaline growth defect, the *rim101*Δ and *sre1*Δ mutants display distinct growth patterns in response to low oxygen availability.

We also tested the sensitivity of the *sre1*Δ and *rim101*Δ mutant strains to cell wall stressors such as Congo Red (inhibits cell wall polymers such as chitin), caffeine (affects cell wall integrity), high salt (osmotically stresses the cell wall), and SDS (stresses the cell membrane) (29). Similar to alkaline pH, high salt resulted in complete growth inhibition for both mutant strains (Fig 1C). In contrast, caffeine did not affect the growth of either mutant (Fig 1C). The *sre1*Δ mutant strain was unable to grow in the presence of Congo Red, whereas the *rim101*Δ mutant strain only showed a subtle growth defect due to this chitin polymer inhibitor (Fig 1C). Also, SDS completely inhibited growth of the *sre1*Δ strain, whereas the *rim101*Δ appeared to be hyper-resistant to the membrane targeting effects of SDS, as evident in the more robust growth of this strain compared to wildtype (29, 30) (Fig 1C).

Sensitivities of mutant strains to cell surface stressors can indicate alterations in the cell wall structure and/or integrity. In addition to providing a protective barrier for the cell, the cell wall serves as an anchor for the attachment of the polysaccharide capsule that can further protect the fungal cells during a human infection (31). The *rim101*Δ mutant strain is known to have a disorganized cell wall and thus a decrease in attached capsule (32, 33). In contrast, the *sre1*Δ mutant strain revealed intact capsule formation (34) (Fig 1E). Overall, these phenotypic comparisons distinguish the *rim101*Δ mutant from the *sre1*Δ mutant in the distinct responses of these strains to cell wall and membrane stress.

### Independent signaling of the Rim and sterol homeostasis pathways

To determine whether the *C. neoformans* sterol homeostasis pathway is specifically activated in response to alkaline pH, we assessed the pH-dependence of the proteolytic cleavage of the Sre1 transcription factor, a marker of pathway activation (21, 23–25, 34). At pH 5.5, the GFP-Sre1 fusion protein remains uncleaved in a 140 kD form (Fig 2A). In contrast, incubating this strain in the same growth medium buffered to pH 8 results in GFP-Sre1 protein cleavage to a 90 kD form, similar to its proteolytic activation in response to hypoxia (Fig 2A and (21)). There was no defect in Sre1 cleavage in the *rim101Δ* mutant strain background (Fig 2B). Therefore, the *C. neoformans* sterol homeostasis pathway is specifically activated by an alkaline pH signal and in a manner that is independent of the Rim alkaline response pathway.

**Figure 2.**
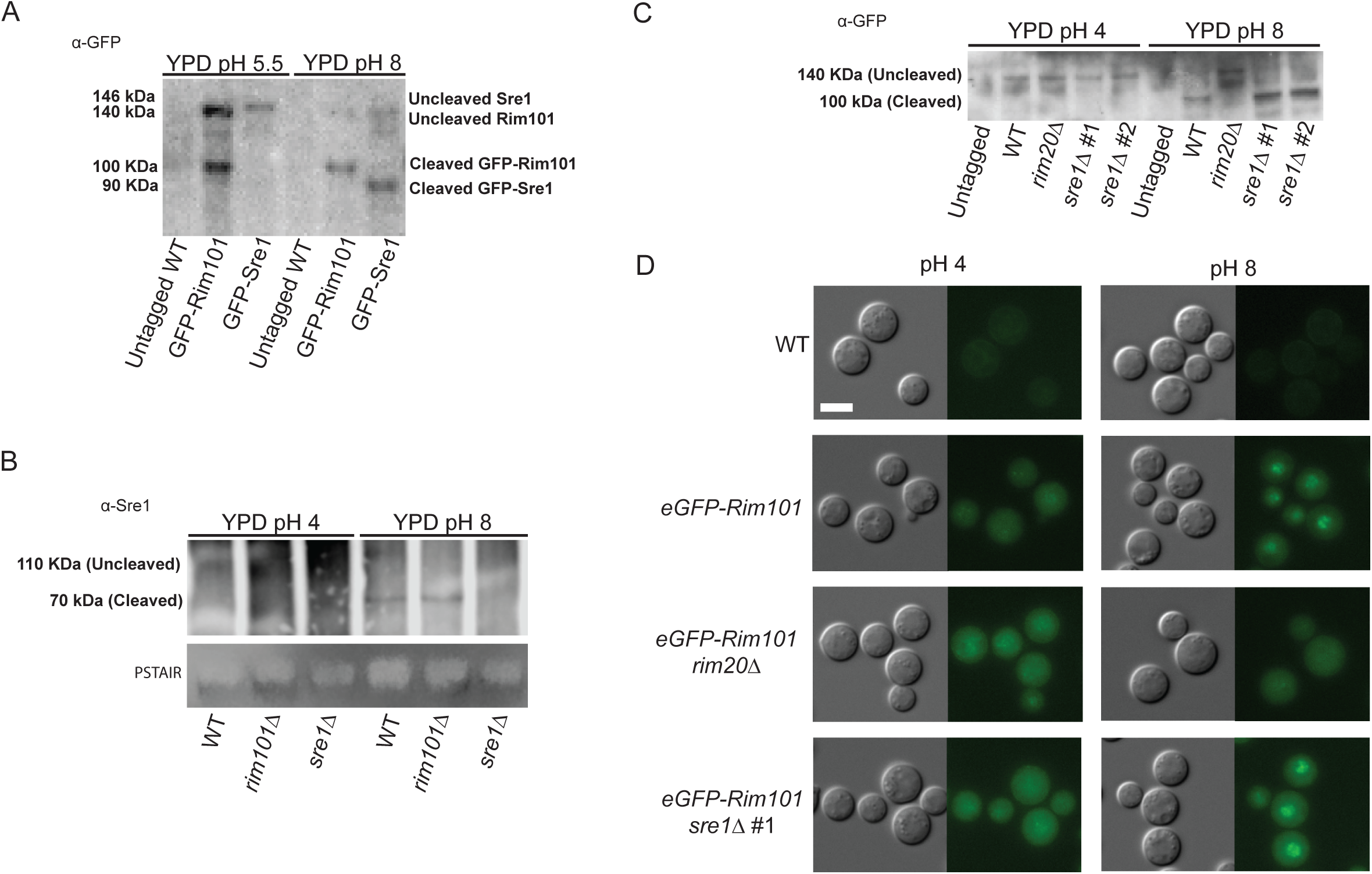
Sre1 activation is dependent on alkaline pH, but not Rim Signaling. **A.** Western blot of both Sre1 and GFP-Rim101 protein processing in low pH and high pH growth conditions. The GFP-Rim101 fusion protein is cleaved from its 140 kDa form to its active 100 kDa form at pH 8. Similarly, the GFP-Sre1 fusion protein is proteolytically processed from 146 kDa to approximately 90 kDa in response to alkaline pH. Indicated strains were incubated for 60-minutes in either pH 5.5 or pH 8 YPD medium prior to lysing. Rim101 and Sre1 protein processing was determined using a GFP-trap pull down and western blotting using an α-GFP antibody. Protein levels were normalized prior to loading. B. Western blot analysis of the Sre1 protein in different genetic backgrounds revealed the cleavage and processing (from 110 kDa to approximately 90 kDa) of the Sre1 transcription factor in the WT and *rim101*Δ mutant backgrounds. Indicated strains were incubated for 60-minutes in either pH 4 or pH 8 YPD medium prior to lysing. Protein processing was determined through Protein-A pull down and western blotting using a polyclonal α-Sre1 antibody. Total protein levels are represented by a PSTAIR loading control. C. The Sre1 protein is cleaved in response to alkaline pH. The *eGFP-RIM101* allele was expressed in the WT, *rim20*Δ mutant and two independent *sre1*Δ mutant strains (*sre1*Δ #1 and *sre1*Δ #2). The untagged WT strain and the eGFP-Rim101 expressing strains were incubated in YPD medium pH 4 or pH 8 for 60 minutes. Rim101 processing was assessed using a GFP-trap pull down and western blotting using an α-GFP antibody. Protein levels were normalized prior to loading. D. The indicated strains (the same as Fig 2C) were incubated in Synthetic Complete media buffered to pH 4 or pH 8 for 60 minutes. Rim101 localization was assessed by epifluorescence microscopy and alkaline-induced nuclear localization was compared to the eGFP-Rim101 positive control. White scale bars indicate 5 microns.

To further define the interaction between the Sre1 and Rim101 signaling pathways, we assessed whether the Sre1 transcription factor is necessary for activation of the Rim pathway as measured by the pH-dependent proteolytic processing and subcellular localization of the Rim101 transcription factor (35). In both the wildtype and *sre1*Δ mutant strains, we observed intact Rim101 processing and cleavage at elevated pH (Fig 2C). Similarly, GFP-Rim101 nuclear localization was enhanced at activating pH in both strain backgrounds (Fig 2D). In contrast, we confirmed both defective protein cleavage and impaired nuclear localization of the Rim101 transcription factor in the *rim20*Δ mutant, a strain lacking a known upstream Rim signaling component (Fig 2C and 2D). These data indicate that the sterol homeostasis pathway is not required for Rim pathway activation.

### The cell wall organization of *sre1*Δ and its *in vitro* immune phenotypes

The *sre1*Δ mutant strain is avirulent in a mouse model of *C. neoformans* infection (25, 34), whereas the *rim101*Δ mutant strain and other Rim pathway mutants have paradoxical hypervirulent phenotypes in the same model (32). In previous work, we demonstrated by transmission electron microscopy that the *rim101*Δ mutant has an aberrant, thick, and disorganized cell wall in comparison to wildtype cells (32). We probed the *rim101*Δ and *sre1*Δ mutant strains with calcofluor white (CFW) and wheat germ agglutinin (WGA) to assess total and exposed levels of chitin, respectively. In both mutant strains, we noted similar increases in cell wall chitin levels as measured by CFW staining. However, the level of exposed chitin (WGA) was only increased in the *rim101Δ* strain. The intensity of the WGA fluorescence was quantified by measuring brightness intensity (FIJI) in photomicrographs (Fig 3A) as well as by flow cytometry (data not shown). The observed increase in total chitin levels can be a non-specific response to cell stress (36). However, increased chitin exposure, as assessed by intensity of WGA staining, has been previously demonstrated to correlate with the degree of macrophage activation *in vitro* (32, 37). Together these cell wall analyses suggest that the Rim pathway and sterol homeostasis pathway induce distinct microbial physiological responses to host-like conditions. Specifically, the Sre1-mediated response to host stress does not include increased exposure of chitin.

**Figure 3.**
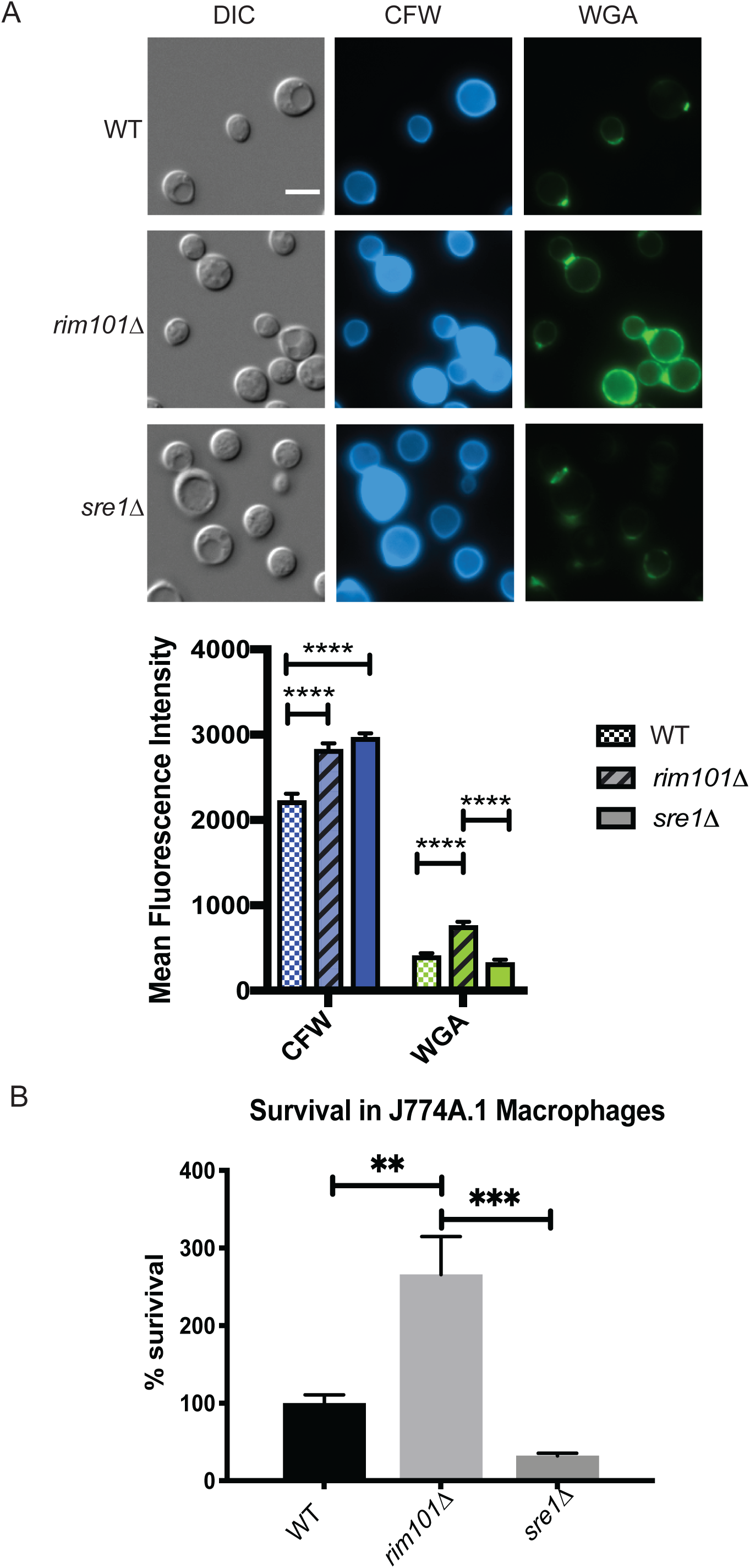
*sre1*Δ and *rim101*Δ mutant strains have varied changes in cell wall chitin exposure and interactions with host immune cells. **A.** Staining of *rim101*Δ, *sre1*Δ and wildtype cells with Calcofluor White (CFW) and Wheat germ agglutinin (WGA). Cells were incubated in CO_2_-independent media for 18 hours at 37°C. Cells were stained with FITC-conjugated WGA and CFW and incubated in the dark for 35 mins and 10 mins respectively. Mean fluorescence intensity was quantified for each strain and each condition. Two-way ANOVA and Tukey’s multiple comparisons test were run to determine statistical significance. White scale bars indicate 5 microns. **** = p-value < 0.0001. B. When grown in the presence of J774A.1 macrophages, the *rim101*Δ mutant strain can survive significantly better than both the wildtype and the *sre1*Δ mutant strain. Indicated strains were co-incubated with macrophages for 24 hours and survival was determined by quantitative cultures. One-way ANOVA and Tukey’s Multiple Comparison tests were run to assess statistical significance between fungal cell survival percentages. 6 biological replicates of each strain were analyzed. ** = p-value < 0.003, *** = p-value < .0002.

To further define the extent to which these cell wall epitopes may affect virulence, we assessed macrophage interactions with the *sre1*Δ mutant compared to the *rim101*Δ mutant strain. Macrophages are among the first immune cells encountered by this pathogen when infecting its host in the human lung. We therefore quantified fungal survival after co-culturing stimulated J774A.1 murine macrophage-like cells with the wildtype, *rim101*Δ, and *sre1*Δ strains. Following co-incubation with macrophages, the *rim101*Δ mutant strain displayed increased survival compared to wildtype, as has been shown previously (33) (Fig 3B). The *sre1*Δ mutant strain displayed a moderate, reproducible reduction in viability in the presence of macrophages compared to the wildtype strain. This result was consistent with the previously reported attenuated virulence of the *sre1*Δ mutant strain in animal models of infection (25, 34). The significantly different patterns of macrophage interaction of the *sre1*Δ and *rim101*Δ mutant strains further suggest that distinct downstream cellular processes are controlled by these alkaline responsive pathways (Figure 3B).

### Ergosterol biosynthesis is required for growth at alkaline pH in *C. neoformans* and other fungal pathogens

Our data support that the Rim and sterol homeostasis pathways are independent cell signaling pathways that each mediate adaptive responses to alkaline stress. Given the established role of fungal Sre1 orthologs in the regulation of membrane sterol content, we hypothesized that alterations in minor membrane lipids, especially ergosterol, might be involved in the adaptive response to alkaline pH. Previous work in *C. neoformans* sterol homeostasis documented decreased ergosterol levels in the *sre1*Δ mutant strain (Bien, Chang, Nes, Kwon-Chung, & Espenshade, 2009; Chang et al., 2009). The *sre1*Δ alkaline pH-sensitivity was rescued by the addition of exogenous ergosterol to the growth medium in a dose-dependent manner (Fig 4A). Importantly, addition of exogenous sterols did not affect the pH of the growth medium. This observation is similar to prior investigations showing growth rescue of various *S. cerevisiae* ergosterol biosynthesis mutants by supplementation with exogenous ergosterol (38). These data suggest that intact ergosterol induction and homeostasis is specifically required for fungal adaptation to alkaline pH.

**Fig 4.**
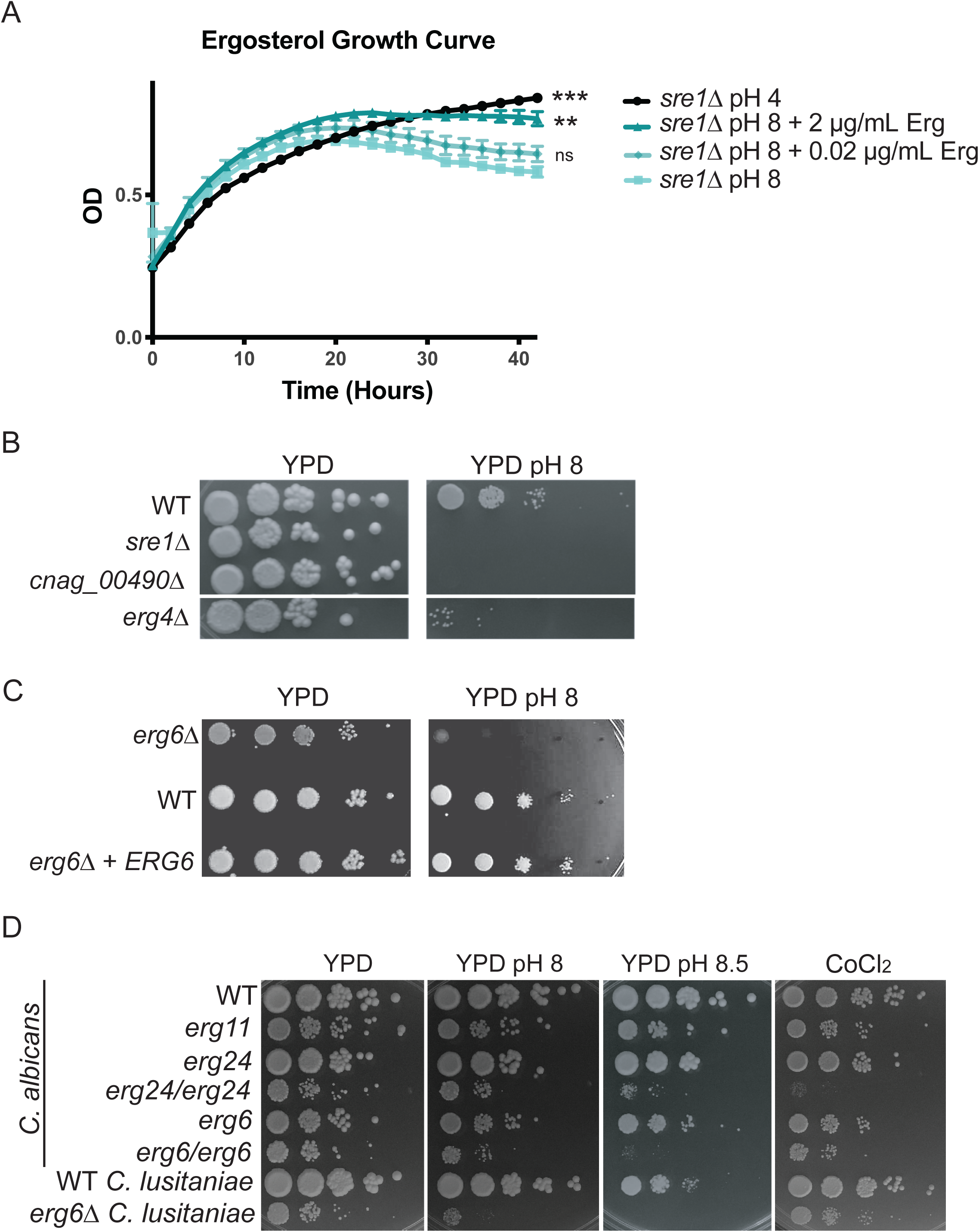
Altered ergosterol content renders strains sensitive to alkaline pH. **A.** The reduced growth rate of the *sre1*Δ mutant strain in liquid growth media at pH 8 can be rescued through the addition of exogenous ergosterol in a dose-dependent manner. Growth rate of indicated strains was assessed by changes in OD_595_ in biological triplicate every 10 minutes for 42 hours at 30°C. Ergosterol was added as indicated. One-way ANOVA and Dunnett’s multiple comparisons test were run on the last time point in each condition compared to the pH 8 alone condition to determine statistical significance. ** = p-value < 0.003, *** = p-value < 0.0005. B. Other sterol-related mutants exhibit alkaline pH-sensitivity. Two deletion mutants related to ergosterol biosynthesis in *C. neoformans* (*erg4Δ* and *cnag_00490*Δ) display a pH-sensitivity when grown on pH 8 growth media. Indicated strains were serially diluted onto YPD medium and YPD 150 mM HEPES pH 8. Growth was compared to WT and assessed after 3 days C. *erg6*Δ *C. neoformans* mutant also exhibits alkaline pH-sensitivity when grown on pH 8 growth. Indicated strains were serially diluted onto YPD medium and YPD 150 mM HEPES pH 8. Growth was compared to WT and reconstituted strains and assessed after 3 days. D. *Candida* species ergosterol mutants reveal similar pH-sensitive phenotypes. *C. albicans* and *C. lusitaniae* wildtype strain and strains with mutations in various components of ergosterol biosynthesis were serially diluted onto YPD medium and YPD 150 mM HEPES pH 8 and 8.5 as well as YES media with 7mM CoCl_2_. Growth was compared to WT and assessed after 2 days.

To further explore the role of ergosterol biosynthesis in the alkaline pH response, we tested three *C. neoformans* ergosterol-related mutants for growth at pH 8, and all shared an alkaline pH growth defect (Fig 4B). Many steps in ergosterol biosynthesis are essential for growth in routine conditions, limiting the availability of *ERG* gene mutants. The non-essential *ERG4* and *ERG6* genes encode terminal enzymes in the ergosterol biosynthesis pathway (22, 39). Compared to wildtype, the *erg4*Δ and *erg6*Δ mutants displayed a specific growth defect at alkaline pH (Fig 4B, 4C and (15)). Similarly, the *CNAG_00490* locus encodes a putative acetyl-CoA acetyltransferase, as does the *ERG10 (CNAG_02918)* gene. The loss-of-function *cnag_00490Δ* mutant also displays alkaline pH-sensitivity (Fig 4B). The pH-sensitivity of the *CNAG_00490* mutant as well as the predictive function of its gene product suggests that it might participate in the conversion of acetyl-CoA to squalene, an early step in sterol synthesis.

Ergosterol is a major component of most fungal membranes, including those of distantly related fungal pathogens in the ascomycete phylum. To further explore the association between sterol homeostasis and alkaline pH response, we tested alkaline pH survival for ergosterol biosynthesis mutants in two *Candida* species: *C. albicans* and *C. lusitaniae*. (Fig 4D). The homozygous diploid *C. albicans erg6*/*erg6* and *erg24/erg24* mutants displayed severe growth defects at high pH that were not evident at more acidic conditions (Fig 4D). Similarly, the haploid *C. lusitaniae erg6* mutant had impaired growth compared to wildtype in alkaline conditions (Fig 4D). These results suggest a conserved requirement for efficient sterol maintenance in the adaptation to alkaline pH among highly divergent fungal species.

### Sre1 regulates membrane-associated transcripts in alkaline growth conditions

The *C. neoformans SRE1*-dependent transcriptome has been defined in the context of the cellular response to low oxygen (21, 22, 34). These prior studies revealed that Sre1 is required for the induction of genes involved in ergosterol homeostasis in an oxygen-dependent manner. However, given the novel role for Sre1 pathway activation at alkaline pH, we defined the pH-responsive Sre1-regulated transcriptional response. Comparison of the transcriptomes of the *sre1*Δ mutant and wildtype after 1.5 hours of growth in alkaline pH revealed 2,655 transcripts that were differentially regulated in a statistically significant manner (adjusted p-value <.05) (Fig S2A and Table S1). This represents approximately one quarter of the *C. neoformans* genome indicating that Sre1 has a major impact on the cell in response to pH stress. Similar to the transcriptome studies in hypoxia, transcript abundance of the majority of the *ERG* genes (13/18) and the *STP1* activating protease was differentially regulated at alkaline pH (Fig 5A and 5B). The *stp1*Δ mutant strain displays a pH-sensitive mutant phenotype similar to the *sre1*Δ mutant strain (15). Importantly, *ERG3* transcript levels had the highest relative fold change in the *sre1*Δ mutant at high pH compared to wildtype (Fig 5A and 5B). *ERG3* encodes a component of the ergosterol biosynthesis pathway and displays similar Sre1-dependent expression in low oxygen conditions (22).

**Fig 5.**
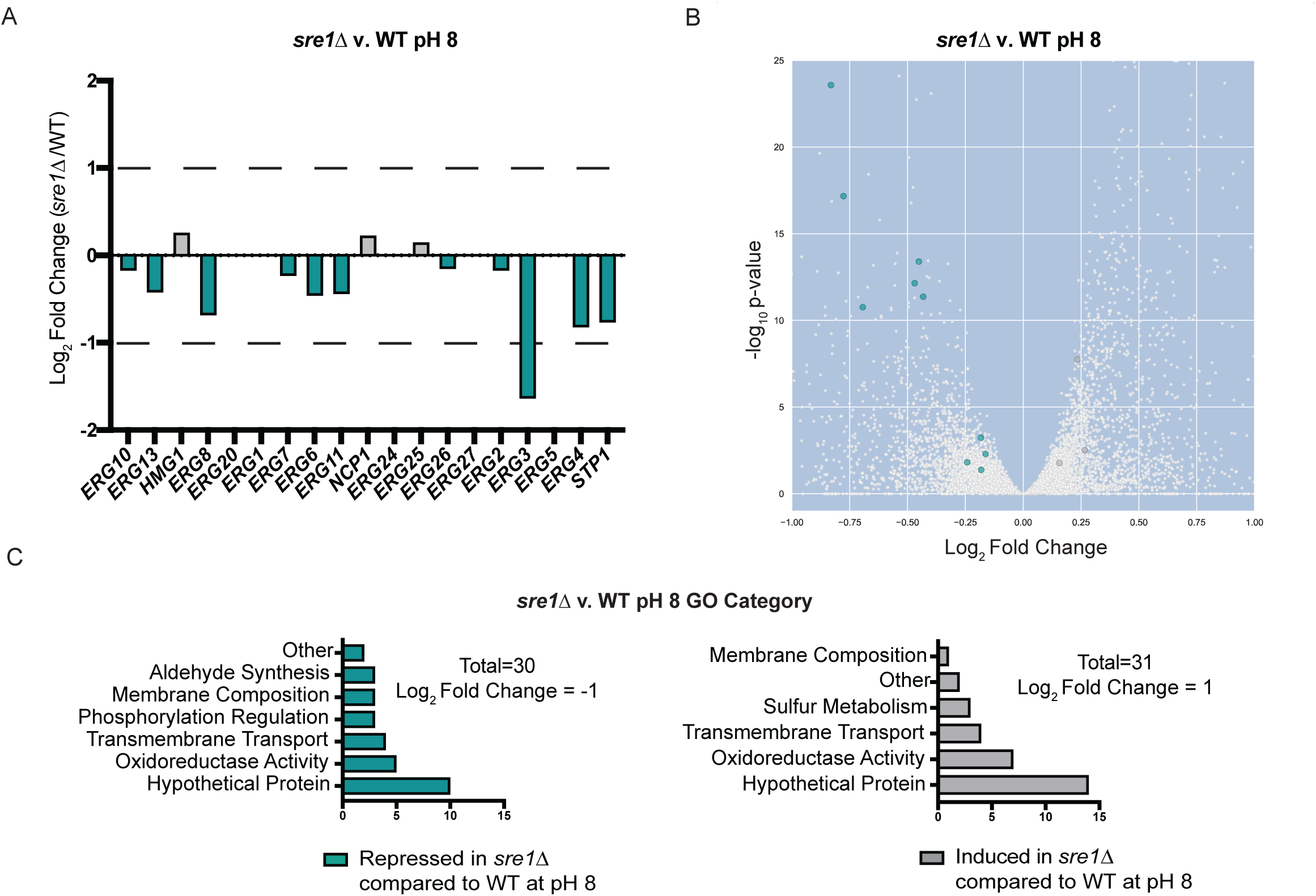
Transcriptomic analysis of the *sre1*Δ and wildtype strains in response to alkaline pH. WT and *sre1*Δ cells were incubated in YPD medium pH 4 or pH 8 for 90 minutes. This experiment was conducted with six biological replicates for each strain and condition. Total RNA was extracted, mRNA was isolated, libraries were prepped and finally sequenced using an Illumina NextSeq 500. GO-term analysis was performed using FungiDB. A. The majority of the known genes in the *C. neoformans* ergosterol biosynthesis were significantly differentially expressed in the *sre1*Δ v. wildtype transcriptome at pH 8. *ERG* genes that were significantly differentially expressed have an adjusted p-value < 0.016 (teal = repressed in the *sre1*Δ mutant compared to wildtype, grey = induced in the *sre1*Δ mutant compared to wildtype). B. Volcano Plot displaying the significantly regulated transcripts in the *sre1*Δ v. wildtype transcriptome at pH 8 (adjusted p-value < 0.05). teal = repressed in the *sre1*Δ mutant compared to wildtype, grey = induced in the *sre1*Δ mutant compared to wildtype). Full volcano plot (zoomed out) in Figure S2. C. GO-term analysis of the *sre1*Δ v. wildtype differentially expressed genes following a 90-minute shift from YPD pH 4 to YPD pH 8. These transcripts were selected based on a strict cutoff of log_2_ fold change = +/-1. Biological processes repressed in *sre1*Δ compared to wildtype at high pH (teal). Biological processes induced in *sre1*Δ compared to wildtype (grey).

**Table 1.**
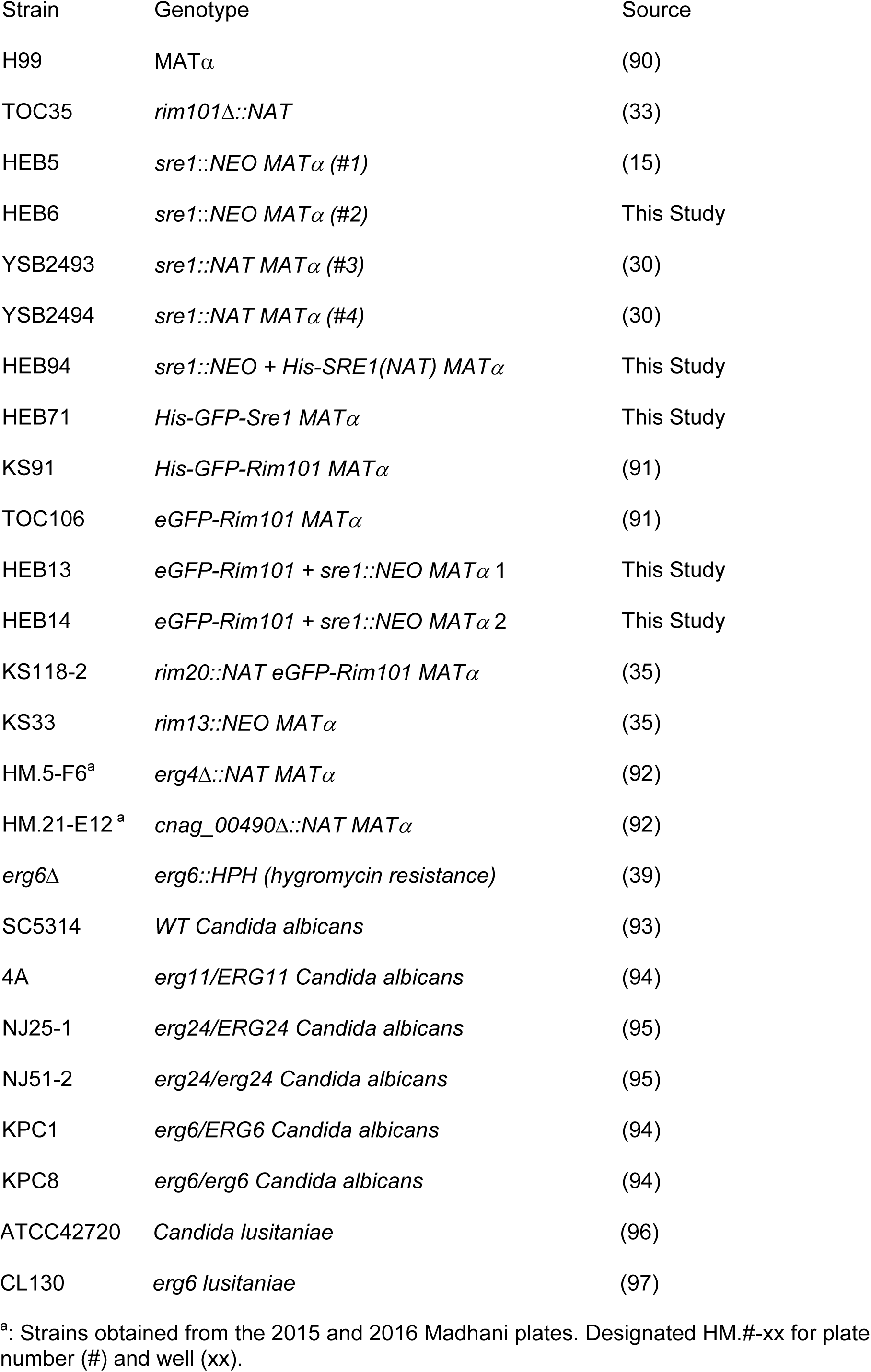
Strain List.

| Strain | Genotype | Source |
| --- | --- | --- |
| H99 | MATα | (90) |
| TOC35 | *rim101*Δ::NAT | (33) |
| HEB5 | *sre1*::NEO MATα (#1) | (15) |
| HEB6 | *sre1*::NEO MATα (#2) | This Study |
| YSB2493 | *sre1*::NAT MATα (#3) | (30) |
| YSB2494 | *sre1*::NAT MATα (#4) | (30) |
| HEB94 | *sre1*::NEO + His-SRE1(NAT) MATα | This Study |
| HEB71 | His-GFP-Sre1 MATα | This Study |
| KS91 | His-GFP-Rim101 MATα | (91) |
| TOC106 | eGFP-Rim101 MATα | (91) |
| HEB13 | eGFP-Rim101 + *sre1*::NEO MATα 1 | This Study |
| HEB14 | eGFP-Rim101 + *sre1*::NEO MATα 2 | This Study |
| KS118-2 | *rim20*::NAT eGFP-Rim101 MATα | (35) |
| KS33 | *rim13*::NEO MATα | (35) |
| HM.5-F6<sup>a</sup> | *erg4*Δ::NAT MATα | (92) |
| HM.21-E12<sup>a</sup> | *cnag_00490*Δ::NAT MATα | (92) |
| *erg6*Δ | *erg6*::HPH (hygromycin resistance) | (39) |
| SC5314 | WT *Candida albicans* | (93) |
| 4A | *erg11*/ERG11 *Candida albicans* | (94) |
| NJ25-1 | *erg24*/ERG24 *Candida albicans* | (95) |
| NJ51-2 | *erg24/erg24* *Candida albicans* | (95) |
| KPC1 | *erg6*/ERG6 *Candida albicans* | (94) |
| KPC8 | *erg6/erg6* *Candida albicans* | (94) |

<sup>a</sup>: Strains obtained from the 2015 and 2016 Madhani plates. Designated HM.#-xx for plate number (#) and well (xx).

Due to the large number of differentially expressed transcripts identified in this analysis, we performed a modified GO-term analysis using FungiDB on genes with a 2-fold or greater change in transcript abundance in the *sre1*Δ mutant compared to wild-type (40). Genes repressed in the *sre1*Δ mutant at high pH are enriched for biological processes such as aldehyde synthesis, cellular respiration/oxidoreduction, membrane composition, phosphorylation regulation, and transmembrane transport. Genes that are induced in this mutant background in alkaline conditions are involved in cellular respiration/oxidoreduction, membrane composition, sulfur metabolism, and transmembrane transport (Fig 5C and Table S1). Interestingly, although some of these GO-terms are shared with the previously published *SRE1* transcriptome in 3% oxygen conditions, the majority of the Sre1-dependent transcripts differ between the two experimental inducing condition: hypoxia versus alkaline pH (22) (Fig S3). Using the same fold-change values to compare these transcript datasets, only nine genes are induced in both conditions, the majority of which are related to ergosterol biosynthesis: *SRE1*, *ERG3*, *ERG11*, *ERG6*, *ERG4*, and *ERG13* (Fig S3). This transcriptome analysis supports the central role for ergosterol biosynthesis genes as potential Sre1-dependent effectors of both hypoxia and the response to alkaline pH. We also documented that different inducing conditions mediate distinct Sre1-dependent transcriptional responses.

We were also able to define groups of genes in the wildtype strain that are either induced and repressed following the shift from low to high pH. These groups include a significant portion of membrane-associated transcripts including integral membrane components, composition regulators, and membrane transporters (Fig S2C and S2D and Table S1). Transcripts with increased abundance in response to alkaline pH include many of the known Rim pathway regulators (*RIM101* and *RIM23*) and pathway outputs (*ENA1*, *CIG1*, and *SKN1*) (Fig S2C and Table S1). Furthermore, many genes involved in membrane composition, glucose/complex carbohydrate metabolism, and regulation of protein phosphorylation were induced in alkaline conditions (Fig S2C). Complex carbohydrates are major components of the fungal cell wall, supporting previous findings that the Rim-mediated pH response is linked to the reorganization of the cell wall (32). GO-term analysis of transcripts with reduced abundance at high pH revealed genes involved in membrane transport, potentially in an effort to regulate import of extracellular ions into the cell (Fig S2D). This analysis revealed no clear repression of membrane composition transcripts at high pH (Fig S2D and Table S1).

### pH affects efficacy of membrane targeting antifungals

Given our observation of a correlation between fungal sterols and growth at alkaline pH, we tested the pH-dependent efficacy of antifungal agents targeting different aspects of membrane ergosterol homeostasis. Amphotericin B (AMB) is a polyene antifungal that removes ergosterol from fungal membranes (41). We observed a dramatic reduction in the AMB minimum inhibitory concentration (MIC) for wildtype *C. neoformans* cells grown on YPD pH 8 (0.25 μg/mL) compared to YPD pH 5.5 (2 μg/mL) (Fig 6A). Furthermore, the time-dependent killing of fungal cells by AMB increased in a pH-dependent manner, further supporting that this drug has a higher efficacy in alkaline growth conditions (Fig 6B). We also found that AMB was significantly more efficacious against the *sre1*Δ strain (MIC = 0.00125 μM) compared to wildtype when the cells were grown at low pH (pH 4-6) (data not shown). The significant increase in AMB activity against this mutant strain with reduced ergosterol content is consistent with our model that disruption in fungal sterols leads to pH-sensitivity. Furthermore, in a drug disc diffusion assay using Pyrifenox, a drug used to treat phytopathogens through inhibition of ergosterol biosynthesis (42), there was a significantly greater zone of clearance and inhibition of growth of wildtype *C. neoformans* cells when grown on media buffered to pH 8 compared to pH 5.5 (Fig 6A).

**Fig 6.**
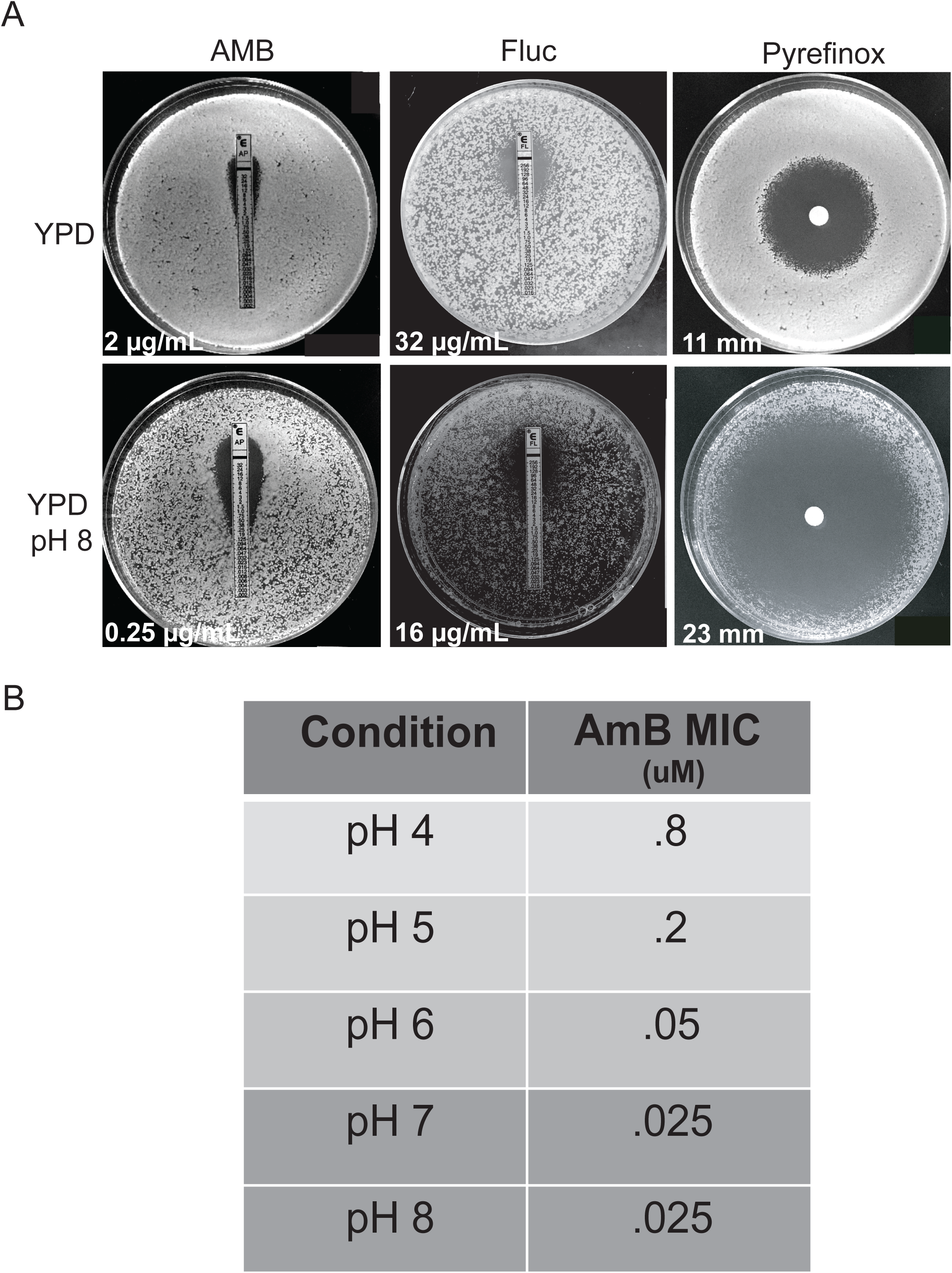
Membrane-targeting antifungals are more active at alkaline pH. **A.** Assessing minimum inhibitory concentrations and the zones of inhibition (white values) of membrane targeting drugs (Amphotericin B (AMB), Fluconazole (Fluc), and Pyrifenox) on wildtype cells grown on YPD or alkaline (YPD pH 8) media. Measurements were taken after 5 days of growth for AMB and pyrifenox and 3 days of growth with Fluc. All plates were incubated at 30°C. B. Minimum inhibitory concentration (MIC) of AMB on wildtype *C. neoformans* grown in increasingly alkaline conditions. MIC was determined after 48 hours of growth at 30°C by broth microdilution.

Fluconazole is an antifungal that inhibits the activity of Erg11, an important component of the ergosterol biosynthesis pathway. We hypothesized that removing ergosterol from the cell membrane in this way would cause a similar sensitivity to alkaline pH that we observed with the ergosterol mutant strains in various fungal pathogens (Fig 4). In contrast to the major pH-dependent activity of AMB and Pyrifenox, we observed a reproducible, but more subtle effect of pH on fluconazole efficacy. The fluconazole MIC was two-fold lower for wildtype at pH 8 (16 μM) compared to YPD pH 5.5 (32 μM) (Fig 6A). The azoles and polyenes have been shown in other organisms, such as *A. fumigatus,* to have variable activity against invasive fungal infections depending on the pH of the growth environment (43). Similar to the findings in *A. fumigatus*, these data support that increases in alkalinity allow for higher efficacy of specific polyenes and azoles against *C. neoformans.* Our data reveal that reduction of ergosterol, either genetically or biochemically using known antifungals, leads to reduced growth in alkaline environments. Altogether, these results further inform the connection between fungal plasma membrane homeostasis, the molecular interactions that drive environment-sensing, and the ability for a biologically diverse group of fungi to grow in increasingly alkaline environments, including their human host.

### Discussion

#### Novel, Rim-independent pH-sensing pathway in C. neoformans

These experiments support a model in which several cell processes and signaling pathways work together to allow microbial growth under stress conditions such as elevated pH. The Rim signaling pathway has been identified in multiple fungal species including *C. neoformans, C. albicans, and S. cerevisiae* as a major signaling response to increases in extracellular pH (1, 2, 47, 48, 3, 15, 32, 33, 35, 44–46) (Fig 7). Its primary function appears to be translating extracellular alkaline pH signals to control adaptive changes in the fungal cell wall ((32) Fig 7B). Data presented in this study identified the sterol homeostasis pathway as a unique mechanism that responds to alkaline pH in a Rim-independent way (Fig 7).

**Fig 7.**
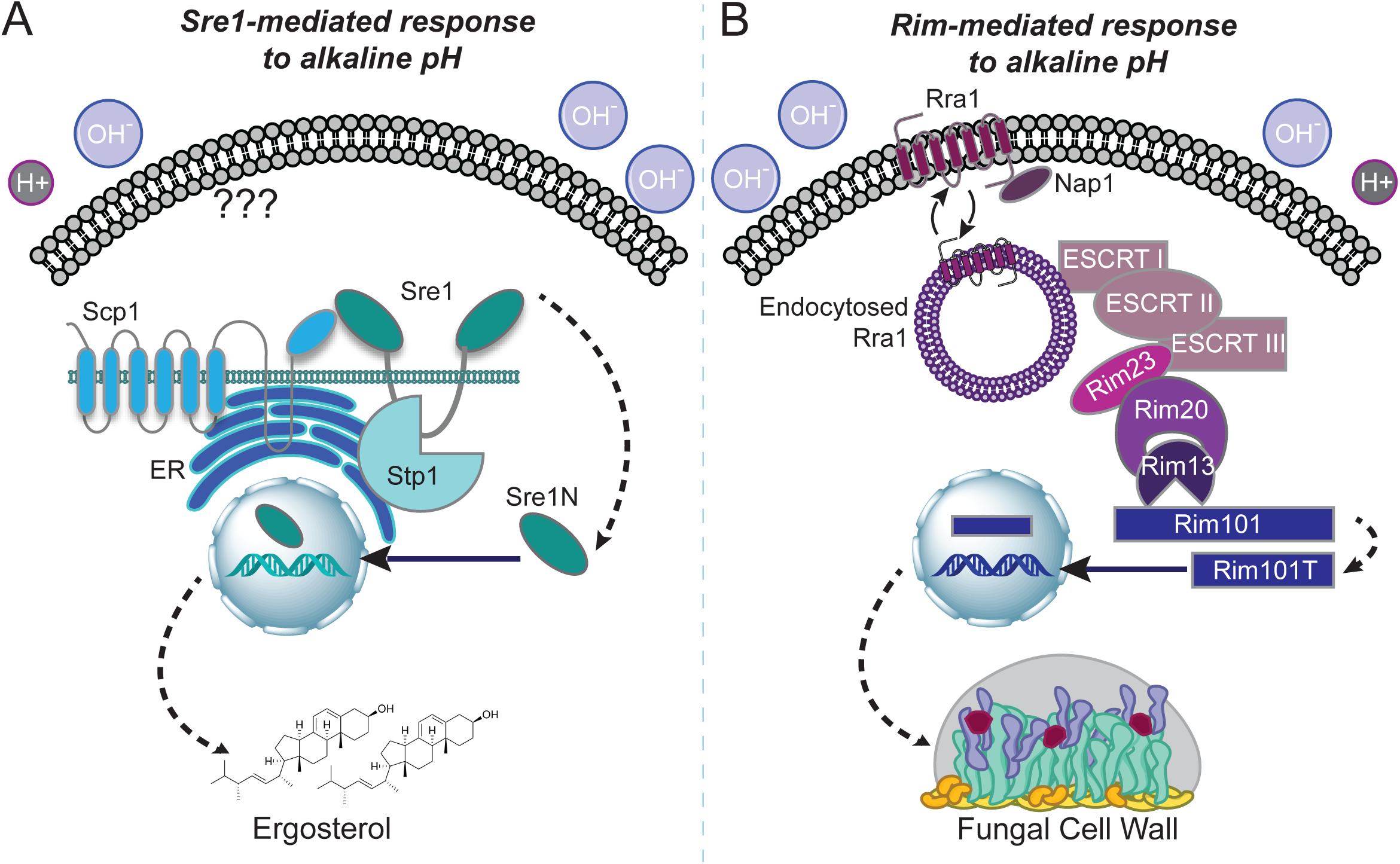
Model of the Sre1-mediated and Rim-mediated distinct responses to physiological pH. A) The activating sensor for the sterol homeostasis pathway is unknown. In response to alkaline pH, the Sre1 transcription factor is cleaved, activated, and localized to the nucleus to aid in the transcription of many genes involved in ergosterol biosynthesis and membrane homeostasis. This cleavage and activation are dependent on both the conserved transmembrane protein, Scp1, and the basidiomycete specific protease, Stp1. B) The Rim alkaline response pathway is signaled through the transmembrane pH-sensor, Rra1. Rra1 is then endocytosed, allowing it to interact with the downstream components of the pathway and propagate the signal to the endosomal membrane complex (ESCRT components, Rim23, and Rim20) and activate the Rim13 protease. This protease cleaves the Rim101 transcription factor allowing it to translocate to the nucleus and induce the expression of genes involved in cell wall and surface remodeling.

The sterol homeostasis pathway has been implicated in the response to alterations in both oxygen availability and membrane ergosterol levels in diverse fungal species. In the fission yeast, *Schizosaccharomyces pombe*, the induction of ergosterol biosynthesis genes by the Sre1 transcription factor and its chaperone proteins (Scp1 and Ins1) has been well characterized in response to hypoxia (18, 20, 49). *C. neoformans*, similar to *S. pombe*, has a well characterized Sre1-mediated response to hypoxia that results in the induction of ergosterol biosynthesis genes to maintain membrane homeostasis (21, 25, 27, 34, 50). However, in *C. neoformans*, a basidiomycete-specific protease has been identified that specifically activates Sre1 in response to hypoxia (Figure 7a) (22, 34). Elements of this pathway have also been identified in the filamentous fungal pathogen *A. fumigatus.* The Sre1 homolog, SrbA, is essential for the ability for this pathogen to grow in environments with limited oxygen (19, 23, 51, 52). This hypoxic response is required for survival in the infected host in which hypoxic microenvironments exist, especially in poorly viable tissue such as necrotic tumors and wounds (53). The dimorphic fungal pathogen, *Histoplasma capsulatum*, also contains a homolog of Sre1 (Srb1) that is essential for the response to hypoxia as well as for virulence (54, 55). Other yeasts such as *C. albicans* and *S. cerevisiae* do not contain genes in their sequenced genomes encoding obvious SREBP homologs. Instead, these species respond to hypoxic stress through the activation of a different transcription factor, Upc2, that directs the induction of ergosterol biosynthesis genes (56, 57). However, the *C. albicans* Cph2 protein binds SRE1-like elements in the genome, and it may therefore be a functional ortholog of Sre1 (58).

The identification of a new role for the sterol homeostasis pathway is informative to better conceptualize and target fungal pathogenesis in general, and cryptococcal pathogenesis in particular for several reasons. First, the sterol pathway in *C. neoformans* has a basidiomycete-specific Stp1 protease that is required for cleavage and activation of Sre1 (22, 25, 34). Genes encoding a similar protease are found in the genomes of other basidiomycete fungi such as *Cryptococcus gattii, Malassezia globosa*, and *Mucor circinelloides* (40), and not in those of more distantly related fungi or higher eukaryotes. This fungal specificity and distinction from the mammalian sterol homeostasis pathway (16, 59, 60) may provide an interesting future target for novel antifungals. Secondly, understanding the extracellular cues that activate this pathway may elucidate more detailed signaling mechanisms controlling sterol homeostasis, potentially revealing some currently unknown upstream components. Presently, it is not known if a common signal in hypoxia or alkaline pH initiates Sre1 signaling, or if multiple upstream Sre1 activators are present (Figure 7a). The *C. neoformans* sterol homeostasis pathway is lacking an obvious INSIG homolog as well as a site-1 protease (24). Elucidating the Sre1-mediated response to alkaline pH through further analysis of our forward genetic screen may uncover either functional orthologs of these proteins or novel pathway components that mediate specific stress responses in *C. neoformans*.

The transcriptional analysis of the *sre1*Δ mutant strain at high pH provided further support for the distinct activation of the Sre1 transcription factor in response to increases in extracellular pH. This type of analysis has been conducted for the *C. neoformans sre1*Δ mutant strain previously, but with conditions of low and high oxygen availability (22). When comparing our transcriptomics data to this previously published microarray analysis, the majority of the transcripts were non-overlapping, suggesting independent downstream effectors of Sre1 in response to specific stress (Fig S3). Furthermore, there was no overlap between the Sre1-associated transcriptome at high pH and the previously published Rim101-associated transcriptome at a similar pH further supporting the distinct nature of these two pH response mechanisms and the specificity of Sre1-mediated response to alkaline pH stress (data not shown and Fig 7).

#### Ergosterol biosynthesis is essential for the ability of fungal pathogens to grow in an alkaline environment

The generation of ergosterol for overall fungal membrane integrity has been well studied in the response to extracellular stresses such as hypoxia and low iron (23, 24, 26, 34, 51, 52). Ergosterol controls the fluidity and structure of fungal cells (61), and it is needed for the formation of microdomains within the membrane containing ion pumps and transmembrane proteins necessary for cellular growth and signaling (12, 13, 61–63). In this study, we have demonstrated that supplementing pH-sensitive mutant strains with ergosterol can rescue the pH-sensitive mutant phenotype, suggesting that the *sre1*Δ mutant pH sensitivity is specifically linked to its ergosterol deficiency.

Our studies further supported this link between ergosterol and the pH response through analysis of the effects of alkaline pH on the biosynthesis of ergosterol at the transcriptional level. In response to a shift in pH, the majority of the known *C. neoformans* ergosterol biosynthesis genes were differentially regulated in the *sre1*Δ strain compared to wildtype. These results support our model and implicate Sre1-mediated membrane homeostasis as a direct response to alkaline stress (Fig 7A). Furthermore, *C. neoformans* and *C. albicans* strains with mutations in known and predicted ergosterol synthetic processes were unable to grow at alkaline pH. These results indicate that ergosterol levels and membrane homeostasis are important in the pH-response mechanisms of many fungal species. This broadens these findings from Sre1-specific regulation of ergosterol affecting pH-growth of a basidiomycete fungal pathogen to general ergosterol maintenance affecting the pH response in many different fungal pathogens across phyla.

#### Ergosterol-depleting antifungals render Cryptococcal cells sensitive to alkaline pH

Our results have not only shown that genetic manipulation of fungal membrane homeostasis and ergosterol biosynthesis can increase the sensitivity of *C. neoformans* to alkaline pH, but also that biochemical and pharmaceutical interventions have the same effect. We tested relevant antifungals that prevent sterol production or directly deplete sterols from fungal membranes and demonstrated that the activity of these drugs improves in neutral/alkaline environments. AMB, an antifungal that directly disrupts the plasma membrane through sequestration of ergosterol (41), was significantly more potent with increases in the pH of the growth environment. Similarly, fluconazole and pyrifenox, drugs that inhibit the ergosterol biosynthesis pathway (42, 64), were also more effective at alkaline pH. These results reflect similar findings in *Aspergillus* species treated with itraconazole and AMB (43). Similar studies using ketoconazole, AMB, and flucytosine (5-FC) against *Candida* species showed that the *in vitro* drug activity increases as a function of pH (65, 66). Interestingly, there has also been one study demonstrating increased efficacy of 5-FC against *C. neoformans* at higher pH (67). The fact that flucytosine does not directly target the cell membrane, together with the subtle alterations in fluconazole activity as a function of pH, suggest that multiple factors control this phenomenon. However, our findings that known ergosterol-targeting antifungals render diverse fungi more vulnerable to growth environments with increasing pH further supports our leading hypothesis that ergosterol homeostasis is a central contributor to the alkaline pH response of many fungal pathogens.

Translating basic investigations in the role of pH modulation in human disease into potential clinical applications has precedent in cancer biology. In mammalian cells, studies of pH regulation in tumor metastasis demonstrated an association between the pH within a tumor and the degree of tumor cell apoptosis, survival, and proliferation (68). The preference among certain malignant cells for more acidic external environments has prompted the exploration of “buffer therapy”, in which site-directed pH modulation is used as an adjunctive therapy to limit tumor growth (69). This type of therapy is also effective against microbial infections that colonize the airways and intestines such as *Pseudomonas aeruginosa* and *Escherichia coli,* respectively (70–72). If these interventions can be used against bacterial infections, one might imagine how similar pH-modulation could be specifically applied to combat the acidic, necrotic core of many established invasive fungal infections, including cryptococcal lesions (53, 73, 74).

Understanding pH-mediated microbial changes in various host micro-niches will allow for the development of optimized antifungal activity at the site of infection.

### Experimental Procedures

#### Strains, media, and growth conditions

Strains generated and/or utilized in this study are shown in Table 1. Each mutant, reconstituted strain, and fluorescent strain was generated in the *C. neoformans* H99 *MATα* genetic background and incubated in either Yeast Peptone Dextrose media (YPD) (1% yeast extract, 2% peptone, and 2% dextrose) or Yeast Nitrogen Base media (YNB). The pH 4, 5, 5.5, 6, 7 and 8 media were made by adding 150 mM HEPES buffer to YPD or YNB media, adjusting the pH with concentrated HCl (for pH < 5.5) or NaOH (for pH > 5.5), prior to autoclaving. Media was supplemented with 20% glucose following autoclaving unless otherwise noted. Cell wall stress phenotypes were assessed by growth on various stress media agar plates as previously described (29). Congo Red (0.5%) and NaCl (1.5M) were added to YPD media prior to autoclaving. Caffeine (1 mg/ml) and SDS (0.03%) were filter sterilized and added to YPD media following autoclaving. Cobalt Chloride plates were made by adding 7 mM (90.89 mg/L) CoCl_2_ solution to autoclaved YES media (glucose, yeast extract, adenine, uracil, histidine, leucine, lysine, and agar) (75, 76). Capsule induction and analyses was completed as previously described (29). Briefly, strains were incubated overnight in YPD media then diluted in tissue culture medium (CO_2_-independent tissue culture medium, TC Gibco) for 72 hours shaking at 37°C, then counterstained with India ink.

The ergosterol supplementation and growth curve analysis was conducted in a 96 well plate. Strains were incubated overnight (∼18 h) at 30°C with 150 rpm shaking. Cells were then pelleted and resuspended in either pH 4 or pH 8 Synthetic Complete media buffered with McIlvaine’s buffer (35). Resuspended strains were added to wells containing the same pH Synthetic Complete media with either 2 μg/ml or 0.02 μg/ml of Ergosterol (Sigma):Tween 80:ethanol (2 mg/ml stock as previously described in (77)). Growth was then measured at an absorbance of 595 nm every 10 minutes for 42 hours, shaking between readings and incubated at 30°C. Control wells containing vehicle alone (ethanol and Tween) were also measured in order to ensure any growth rate change detected was due to the addition of ergosterol. One-way ANOVA and Dunnett’s multiple comparisons test were run on the last time point in each condition compared to the pH 8 alone condition to determine statistical significance. The pH of the media in the wells was tested following the experiment to ensure the media remained buffered.

To generate the *sre1*Δ deletion and *eGFP-Rim101* + *sre1*Δ deletion and tagged deletion constructs, respectively, we performed the previously described double-joint PCR with split drug resistance marker method to make targeted gene deletions (15, 78). In brief, we generated the following two PCR products: 5’ flanking region of the target locus (1000 bp) with a truncated drug resistance cassette and the remainder of the drug resistance cassette with the 3’ flanking region of the target locus (1000 bp). We then used biolistics to transform these two amplicons into either the wild-type *C. neoformans* strain (H99) or the *C. neoformans* strain that contains endogenously expressed GFP-Rim101 (79). Transformants were selected for the presence of the construct on YPD medium + neomycin (NEO). To generate the fluorescently tagged *His-GFP-Sre1* strain, we used In-Fusion (Clontech) to clone the *SRE1* gene and terminator into the HGNAT (pCN19) plasmid, containing the GFP sequence and the nourseothricin (NAT) resistance marker (80). This plasmid was then biolistically transformed into the H99, WT strain. To generate the SRE1 reconstituted strain we cloned the SRE1 gene and terminator into the pCH233 plasmid, containing the nourseothricin (NAT) resistance marker (81). This plasmid was then biolistically transformed into the *sre1*Δ (HEB5) strain. The primers used to generate each strain are listed in Table 2. Primers used to validate all *sre1*Δ mutants through Southern analysis (data not shown) are also listed in Table 2. Transformants were selected on YPD medium containing NAT (fluorescent strain) or NAT/NEO (reconstituted strain). Plasmids used in this study to amplify markers and clone new plasmids are listed in Table 3.

**Table 2.**
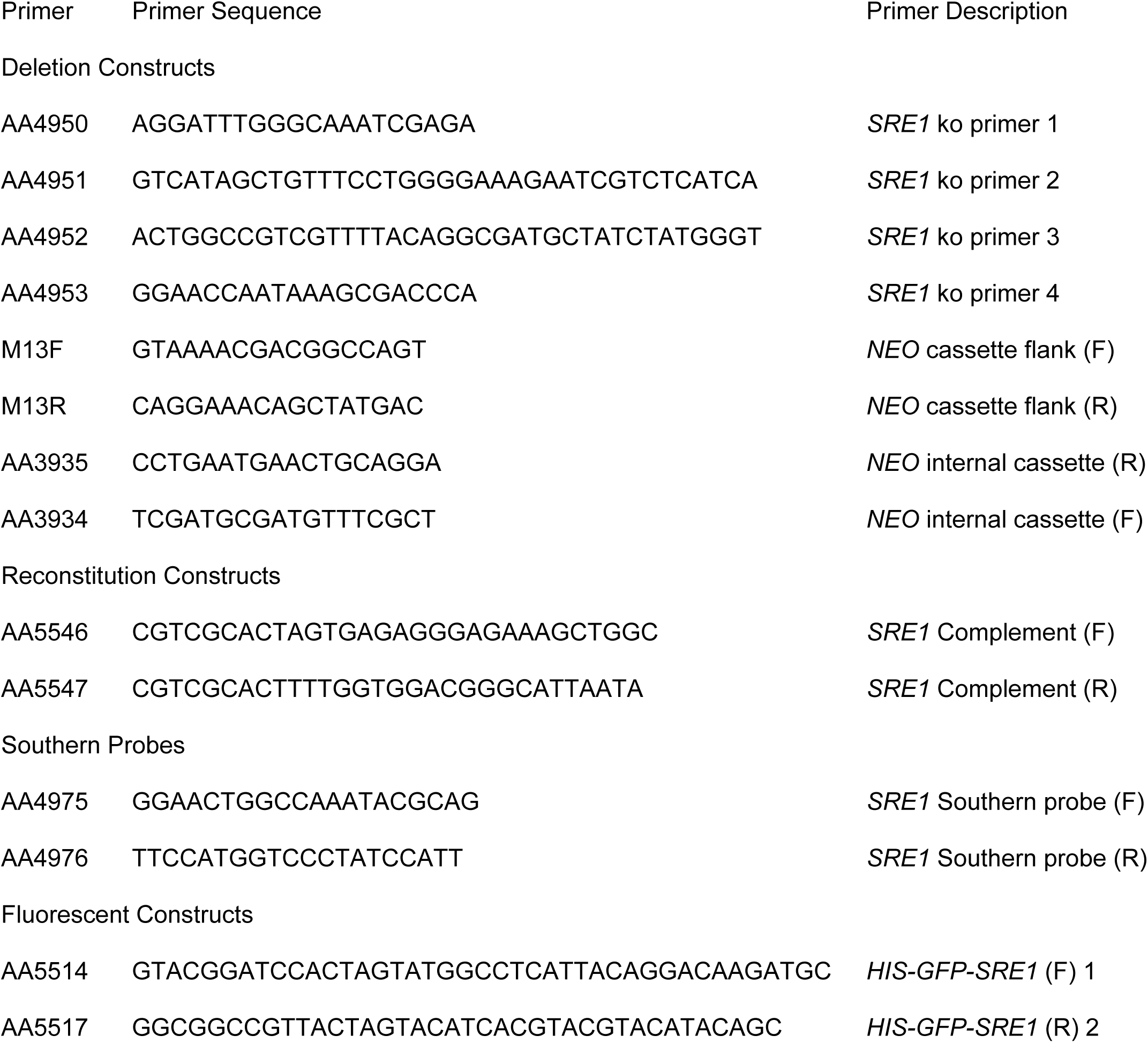
Primers used in this study.

**Table 3.** **Plasmids used in this study**

|  |  |  |  |  |
| --- | --- | --- | --- | --- |
| 641 | Table 3. Plasmids used in this study |  |  |  |
|  | Plasmid | ORF | Backbone | Source |
|  | pJAF | Neomycin (NEO) Resistance Cassette |  | (98) |
|  | pCN19 | Histone H3 promoter; GFP | pJAF | (80) |
|  | pCH233 | Nourseothricin (NAT) resistance cassette |  | (81) |
|  | pHEB13 | Histone H3 promoter; GFP; *SRE1* including terminator | pCN19 | This Study |

#### Microscopy

To analyze GFP-Rim101 localization in the WT, *rim20*Δ and *sre1*Δ backgrounds, strains were incubated overnight (∼18 h) at 30°C with 150 rpm shaking. Cells were then pelleted and resuspended in either pH 4 or pH 8 Synthetic Complete media buffered with McIlvaine’s buffer. Strains were shaken at 150 rpm and 30°C for 60 minutes as this has been shown to be sufficient time to observe the nuclear localization of Rim101 in WT cells (15). Fluorescent images were captured using a Zeiss Axio Imager A1 fluorescence microscope equipped with an Axio-Cam MRM digital camera. Images were created using ImageJ software (Fiji) (82).

#### Protein Extraction, Immunoprecipitation, and Western Blot

Protein extracts were prepared as in a similar manner to what was previously described (15). Briefly, strains were incubated for ∼18 hr at 30 °C with 150 rpm shaking in YPD media buffered to pH 4 or 5.5 with HEPES and HCl. Cells were then pelleted and resuspended in YPD media buffered to pH 8 with HEPES and NaOH. These cells were incubated for 60 minutes and immediately pelleted and flash frozen on dry ice. Lysis was performed by bead beating (0.5 ml of 3 μM glass beads in a Mini-BeadBeater-16 (BioSpec) for 6 cycles of 30 seconds each with a one-minute ice incubation between bead-beating cycle for cell recovery). Supernatants were washed 3 times with 0.4 ml of lysis buffer (2x protease inhibitors (Complete, Mini, EDTA-free; Roche), 1x phosphatase inhibitors (PhosStop; Roche) and 1 mM phenylmethanesulfonyl-fluoride (PMSF). The crude pellet was pelleted through centrifugation at 15,000 rpm, 4 °C, for 5 minutes, and the supernatant (cell lysate) was transferred (∼ 1 mL) to a new tube. For westerns assessing the presence of protein by probing for GFP, 50 μl of lysate was saved as whole lysate and 25 μl of GFP-trap (Chromotek) resin (equilibrated and resuspended in lysis buffer) was added to the remaining lysate. The lysates containing GFP-trap were incubated at 4°C for 2 hours, rotating. Following incubation, lysates were spun down (2,500 g for 2 minutes at 4°C) and washed 3 times with detergent-free buffer containing 2x protease inhibitors (Complete, Mini, EDTA-free; Roche), 1x phosphatase inhibitors (PhosStop; Roche) and 1 mM phenylmethanesulfonylfluoride- fluoride (PMSF). GFP-trap resin was resuspended in 4X NuPage lithium dodecyl sulfate (LDS) loading buffer and 10X NuPage Reducing Agent. Western blots were performed using a 4-12% NuPage BisTris gel. To probe and detect GFP-Rim101 and GFP-Sre1, immunoblots were incubated with an anti-GFP primary antibody (using a 1/10,000 dilution, Roche), followed by a secondary anti-mouse peroxidase-conjugated secondary antibody (using a 1/25,000 dilution, Jackson Labs). Proteins were detected by enhanced chemiluminescence (ECL Prime Western blotting detection reagent; GE Healthcare).

For western blots assessing the presence of cleaved and uncleaved Sre1 using the polyclonal α-Sre1, lysates were prepped in the same way as previously described. Following lysis and initial centrifugation of the crude pellet, 500 μl of lysates were pre-cleared with 30 μl Protein A-Agarose (Sigma) and rotated for 1 hour at 4°C. Lysates were incubated with 5 μl of α-Sre1 polyclonal antibody (generously given to us by the Espenshade lab (21)) for one hour.

Protein A (60 μl/sample) was washed twice in lysis buffer and resuspended in equal volumes. Equilibrated Protein A was then added to each lysate and incubated at 4°C for 1 hour, rotating. Following incubation, lysates were spun down (2,500 g for 2 minutes at 4°C) and washed 2 times with lysis buffer, 1 time with lysis buffer + 1M NaCl, and 2 times with lysis buffer. Protein A resin was then resuspended in 4X NuPage lithium dodecyl sulfate (LDS) loading buffer and 10X NuPage Reducing Agent. Western blots were performed using a 3-8% NuPage Tris Acetate gel, with Tris Acetate Running buffer. To probe and detect Sre1, immunoblots were incubated in α-Sre1 primary antibody (using a 1/200 dilution, (21)) and then in secondary anti-rabbit peroxidase-conjugated secondary antibody (using a 1/50,000 dilution, Jackson Labs). Proteins were detected in the same way as described above.

#### Cell Wall Staining and Quantification

For chitin and exposed chitin detection, cell wall staining with wheat germ agglutinin (WGA) and calcofluor white (CFW) was assessed as previously described (29). Briefly, overnight YPD cultures were diluted 1:10 in CO_2_-independent liquid medium and incubated (∼18 h) at 37°C with 150 rpm shaking. Cells were stained with 100 μg/ml of FITC-conjugated WGA and 25 μg/ml CFW and incubated in the dark for 35 mins and 10 mins respectively. Quantitative analysis using ImageJ software was performed as previously described (29, 37).

#### Macrophage Survival Assay

J774A.1 cells were incubated in a humidified 37°C incubator with 5% CO_2_, passaged twice weekly, and were kept in tissue culture flasks in 20-25 ml of macrophage medium (DMEM, heat-inactivated fetal bovine serum-FBS, Penicillin-streptomycin (Gibco 15140-122), and MEM non-essential amino acid solution (Gibco 11140-050)). Survival of *C. neoformans* strains within alveolar macrophage-like J744A.1 cells was assessed by aliquoting 100 μl of 10^5^ viable cells into a 96-well plate, avoiding edges as previously described (83). The plates were incubated overnight in 37°C incubator with 5% CO_2_. Macrophages were then activated with 10 nm phorbol myristate acetate (PMA) and incubated at 37°C 5% CO_2_ for 1 hour. Fungal cells were incubated overnight (∼18 h) at 30°C with 150 rpm shaking. Cells were then pelleted, washed twice in PBS, and resuspended in Macrophage medium. Fungal cells (10^6^ cells/ml) are opsonized with mAb 18B7 (1μg/ml) for one hour at 37°C. Cell concentrations were verified with quantitative culture. Macrophage medium was removed from the 96 well plate, and 100 μl of opsonized fungal cells are added to each well. The co-cultures were incubated for 1 hour at 37°C incubator with 5% CO_2_. Each well was then washed 3 times with PBS to remove extracellular yeast. 100 μl of macrophage medium was added to each well and incubated for 24 hours at 37°C with 5% CO_2_. Following incubation, macrophage killing was determined by adding 200 μl sterile dH_2_O to each well, incubating at room temperature for 5 minutes, and assessing by quantitative cultures. One-way ANOVA and Tukey’s Multiple Comparison tests were run to assess statistical significance between fungal cell survival percentages. 6 biological replicates of each strain were analyzed.

#### RNA-sequencing preparation and analyses

WT and *sre1*Δ cells were incubated at 30°C with 150 rpm shaking in YPD media to mid-logarithmic phase. Approximately 1x10^9^ cells from each strain were pelleted and resuspended in YPD media buffered to pH4 or pH8 and incubated at 30°C for 90 minutes with 150 rpm shaking. All cells were pelleted, flash frozen on dry ice, and lyophilized overnight. This experiment was conducted with six biological replicates for the WT strain and the *sre1*Δ strain in both pH4 and pH8 conditions (24 samples total). RNA was isolated using the Qiagen RNeasy Plant Minikit with optional on column DNase digestion (Qiagen, Valencia, CA). RNA quantity and quality were measured using the Agilent 2100 Bioanalyzer. The NEBNext Poly(A) mRNA Magnetic Isolation Module was used to enrich for mRNA and the NEBNext Ultra II Directional RNA Library Prep Kit for Illumina was used to prepare libraries (New England Biolabs, Ipswich, MA).

Libraries were submitted to the Duke Sequencing and Genomic Technologies Shared Resource for sequencing on the Illumina NextSeq 500 with 75 base pair, single-end reads.

Reads were mapped to the *C. neoformans* H99 reference genome (obtained from NCBI, accessed July 2019) using STAR alignment software (84). Differential expression analyses were performed in R using an RNA-Seq Bioconductor workflow (85, 86) followed by the DESeq2 package with a false discovery rate (FDR) of 5% (87). Genes were considered statistically differentially expressed if they had an adjusted p-value < 0.05.

A modified Gene Ontology-term (GO-term) analysis using the FungiDB database was performed to identify genes that were significantly regulated in a given process as previously reported (15, 88). The differentially expressed genes in each category were determined based on two criteria: p-value < .05 and base mean value > 20. Further differentiation was made based on the log_2_ fold change values. For the *sre1*Δ v wildtype dataset, we used a log_2_ fold change = +/-1. For the positively regulated genes in the wildtype pH 4 v pH 8 dataset we used log_2_ fold change = 1 and for the negatively regulated genes in the wildtype dataset we used a log_2_ fold change = -3 due to the large amount of genes in this set. Fold change graphs were generated in GraphPad Prism (GraphPad Prism version 8.00 for Mac, GraphPad Software, San Diego California USA, www.graphpad.com) and Seaborn was used to visualize the DESeq2 results in a Volcano Plot (89). A complete list of the RNA-Seq datasets containing differentially expressed genes in each strain and associated with the appropriate GO term category can be found in Table S1.

#### Antifungal Susceptibility Tests

Fluconazole and Amphotericin B (AMB) E-Test assays and Pyrifenox disc diffusion: Fungal cells were incubated overnight (∼18 h) at 30°C with 150 rpm shaking in YPD. Cells were normalized to an OD_600_= 0.6 and diluted 1:10 in PBS and 100 μl were plated to either YPD or YPD pH 8 agarose plates. For the Fluconazole and AMB E-test assay, an E-test strip (Biomerieux) containing a gradient of drug concentrations was placed on top of the plated fungal lawn. Plates were then incubated at 30°C for 72 (AMB) and 120 (Fluconazole) hours. Pyrifenox susceptibility was assessed by standard disc diffusion assays using 5 μl Pyrifenox (Sigma-Aldrich CAS Number: 88283-41-4, final concentration of 1.2 g/mL). Plates were then incubated at 30°C for 72 hours. Zones of inhibition were determined as a surrogate of antifungal activity.

Minimum Inhibitory Concentration testing of AMB against a pH gradient was performed by broth microdilution: AMB resuspended in DMSO was serially diluted in Synthetic Complete media buffered to pH 4, 5, 6, 7, or 8 with McIlvaine’s buffer a 96 well plate with the highest concentration = 3.2 μg/ml Fungal cells were incubated overnight (∼18 h) at 30°C with 150 rpm shaking in YPD. Cells were then normalized and diluted in Synthetic Complete media buffered to pH 4, 5, 6, 7, or 8 with McIlvaine’s buffer and added to the corresponding pH well containing Amp B. Plates were incubated at 30°C for 72 hours, and the MIC was determined to be the lowest concentration of drug that led to no fungal cell growth.

#### Data Availability

All raw and analyzed RNA-sequencing data has been submitted to the NCBI GEO database GSE147109. https://www.ncbi.nlm.nih.gov/geo/query/acc.cgi?acc=GSE147109.

## Acknowledgements

We acknowledge the Duke Sequencing and Genomic Technologies Shared Resource for their assistance with the various projects in this study. We thank Josh Granek and the other staff of the High Throughput Sequencing Course held through the Department of Biostatistics and Bioinformatics at Duke University for their assistance with our RNA sequencing library prep and analysis. We thank the Espenshade Laboratory for their generous gifting of the α-Sre1 polyclonal antibody. We also thank the Madhani Laboratory and the NIH funding (R01AI100272) for the *erg4*Δ and *cnag_00490*Δ deletion strains available through the Fungal Genetics Stock Center. We thank Max Moskovitz for his assistance with phenotyping sterol-related deletion strains. We thank the Heitman Laboratory for sharing the various *Candida* deletion strains. Finally, we acknowledge our own NIH funding (R01 AI074677, 1F31A140427-01A1, P01 AI104533) that made these studies possible. The authors have no relevant conflicts of interest.

## Author Contributions

HEB, CLT, JWS and JAA were involved with the conception and design of experiments and the writing process. HEB, CLT, and LF were involved in the acquisition of the data. All authors participated in the analysis and interpretation of the data.

---

**Fig S1. The *sre1*Δ pH-sensitive and hypoxia-sensitive mutant phenotypes can be rescued with the addition of the wildtype *SRE1* allele.** Strains were spotted in serial dilutions onto YPD, YPD pH 8, and YES + 7 mM CoCl_2_. Growth was assessed after 3 days of growth and compared to both the *sre1*Δ mutant and WT strains.

**Fig S2. Full transcriptome analyses and WT GO terms**

A. Volcano plot of the full transcriptome as analyzed from the *sre1*Δ v. WT at pH 8 dataset (cropped version in Fig 5B).

B. Volcano plot of the wildtype transcriptome. GO-term analysis of the wildtype differentially expressed genes that are (C) induced and (D) repressed following a 90-minute shift from YPD pH 4 to YPD pH 8. These transcripts were selected based on a strict cutoff of log_2_foldchange = +1 for induced transcripts log_2_foldchange = -3 for repressed transcripts based on the uneven distribution of total repressed transcripts shown on the volcano plot (B).

**Figure S3. Compared transcriptomics.** Venn’s Diagram showing the overlap between three different transcriptome datasets in *C. neoformans* serotype A (H99). Bien (2009) performed a comparative transcriptomics microarray study using the *sre1*Δ mutant strain and the *sre1*Δ + *SRE1* complemented strain in low oxygen growth environments (3% O_2_) (dark blue). A 1.4 fold-change cutoff was applied to this dataset and the transcripts included in this comparison were increased in the reconstitute compared to the mutant. Our data set looking at the *sre1*Δ mutant compared to wildtype at low (4) and high pH (8) is shown in teal. We originally applied a fold-change cutoff of 2 to our dataset to analyze our GO-term categories (Figure 5c and 5d), but lowered the cut-off to 1.4 to match Bien (2009) and only included those transcripts that were repressed in the *sre1*Δ mutant compared to wildtype. The overlap of our data with the previously published dataset revealed many ergosterol-related transcripts: *SRE1, ERG3*, *ERG4*, *ERG11*, *ERG6*, and *ERG13*.

**Table S1. RNA-seq analysis**

Page 1. *sre1*Δ v. wildtype differentially expressed genes at pH 8 (repressed). Annotation of GO-terms and putative processes/functions included. p < 0.05 and log_2_fold change < -1. Page 2. *sre1*Δ v. wildtype differentially expressed genes at pH 8 (induced). Annotation of GO-terms and putative processes/functions included. p < 0.05 and log_2_fold change < 1. Page 3. *sre1*Δ v. wildtype entire transcriptome at pH 8. Raw data. Page 4. Wildtype pH 4 v. wildtype pH 8 entire transcriptome. Raw data. Page 5. Wildtype pH 4 v. wildtype pH 8 differentially expressed genes (induced). Annotation of GO-terms and putative processes/functions included. p < 0.05 and log_2_fold change < 1. Page 6. Wildtype pH 4 v. wildtype pH 8 differentially expressed genes (repressed). Annotation of GO-terms and putative processes/functions included. p < 0.05 and log_2_fold change < -3

